# Viral inhibition of the anaphase promoting complex enhances replication by elevating nucleotide pools

**DOI:** 10.64898/2026.01.26.701660

**Authors:** Owen J. Chen, Cynthia Y. Feng, Reese Jalal Ladak, Maximilian A. Crosby, Mariana de Sá Tavares Russo, Yilin Wang, Paola Blanchette, Yingke Liang, David M. Sharon, Mohamed Moustafa-Kamal, Isabelle Gamache, Nahum Sonenberg, Daina Avizonis, Jose G. Teodoro

**Author notes:** Corresponding author: (JGT). These authors contributed equally to this work (Owen J. Chen, Cynthia Y. Feng).

## Abstract

The anaphase promoting complex/cyclosome (APC/C) is a large, ubiquitin ligase and a central regulator of cell cycle progression. By targeting key substrates for degradation during mitosis and G1 phase, the APC/C coordinates metabolic fluctuations that occur during the cell cycle. A diverse range of viruses have convergently evolved mechanisms to bind and inhibit the APC/C; however, a molecular understanding for these interactions has never been demonstrated. Here, we use chicken anemia virus (CAV), a small single-stranded DNA virus encoding only three proteins, to demonstrate the importance of viral APC/C inhibition during replication. We show that the Vp3 protein of CAV inhibits the APC/C, causing a dramatic mitotic arrest during infection. Mutant virus lacking Vp3 are defective for replication and can be rescued by APC/C inhibition. Metabolomic profiling during CAV infection revealed that Vp3 expression mediates a broad increase in nucleotide pools. Moreover, viral inhibition of the APC/C resulted in stabilization of enzymes required for nucleotide biosynthesis. These findings suggest that the APC/C is a general target of many viruses to elevate nucleotide levels and facilitate viral genome replication.

**Author Summary:** Viruses dramatically reshape the cells they infect in order to replicate. To do this efficiently, many viruses disrupt the host’s normal control systems, particularly those that regulate cell growth and division. One such system is the anaphase promoting complex/cyclosome (APC/C), a key cellular machine that controls when specific proteins are broken down during the cell cycle. Surprisingly, a wide range of viruses—including adenovirus, HIV, human papillomavirus, herpesvirus, and others—have been shown to interfere with the APC/C. Until now, however, it was unclear why so many unrelated viruses target this same pathway. In our study, we used chicken anemia virus (CAV), one of the smallest known animal viruses, as a simple model to uncover the underlying mechanism. CAV produces just three proteins, one of which—called VP3—strongly inhibits the APC/C. We found that this inhibition increases the availability of nucleotides, the basic building blocks needed to copy viral DNA. By disrupting normal cell cycle control, the virus effectively redirects cellular resources toward its own replication. These findings reveal APC/C inhibition as a common and powerful strategy used by diverse viruses, offering new insight into how viruses exploit host cells and highlighting potential targets for future antiviral therapies.

## Introduction

Viruses induce enormous changes in their host cells during the course of infection to facilitate replication. For example, viruses are well known to dysregulate cell cycle dynamics (1). A diverse range of viruses have been demonstrated to associate with and inhibit a key cell cycle regulator known as the anaphase promoting complex/cyclosome (APC/C). Papillomavirus (2), adenovirus (3), herpesvirus (4), poxvirus (5), retrovirus (6) and anellovirus (7) have all been shown to encode proteins that bind and inactivate the APC/C. The APC/C is a large 1.5-MDa complex composed of at least fifteen subunits and is required for cell division in all eukaryotes (8). The APC/C functions as an E3 ubiquitin ligase that targets specific substrates for degradation via the 26S proteosome at transition points in the cell cycle (8). The APC/C must be bound to a co-activator to have ubiquitylation activity and has different substrate specificities depending upon which co-activator is bound (8). During anaphase, the APC/C is bound to the co-activator CDC20, which is responsible for coordination of mitotic events by targeting substrates such as the mitotic cyclins and securin (8). A second APC/C complex bound to the co-activator CDH1 exists in G1. APC/C^CDH1^ has a unique set of substrates that control S phase entry and DNA replication (9–12). The first characterized substrates of the APC/C were the mitotic A-and B-type cyclins (13). For cell division to occur successfully, a series of events must be perfectly coordinated including DNA synthesis, spindle formation, and nuclear division.

We previously demonstrated that the Vp3 protein encoded by chicken anemia virus (CAV) binds to the APC/C and promotes the disassembly of the complex (7). CAV is a small, single-stranded DNA virus in the *Anelloviridae* family and is among the smallest viruses that infect animals (14). The CAV genome is 2.3 kilobases and encodes only three open reading frames (*VP1*, *2* and *3*, see Fig 1A)(15, 16). Vp3, also referred to as apoptin, is a small 13.6-kDa protein that mediates the pathology of the virus in chickens (17). CAV has a natural tropism for rapidly dividing cells and Vp3 has attracted considerable interest because it induces cell death preferentially in tumour cells (18–22). Vp3 induces mitotic arrest in a manner that is dependent on binding the APC/C (7, 23), but the importance of this function for viral replication has never been addressed.

**Fig 1.**
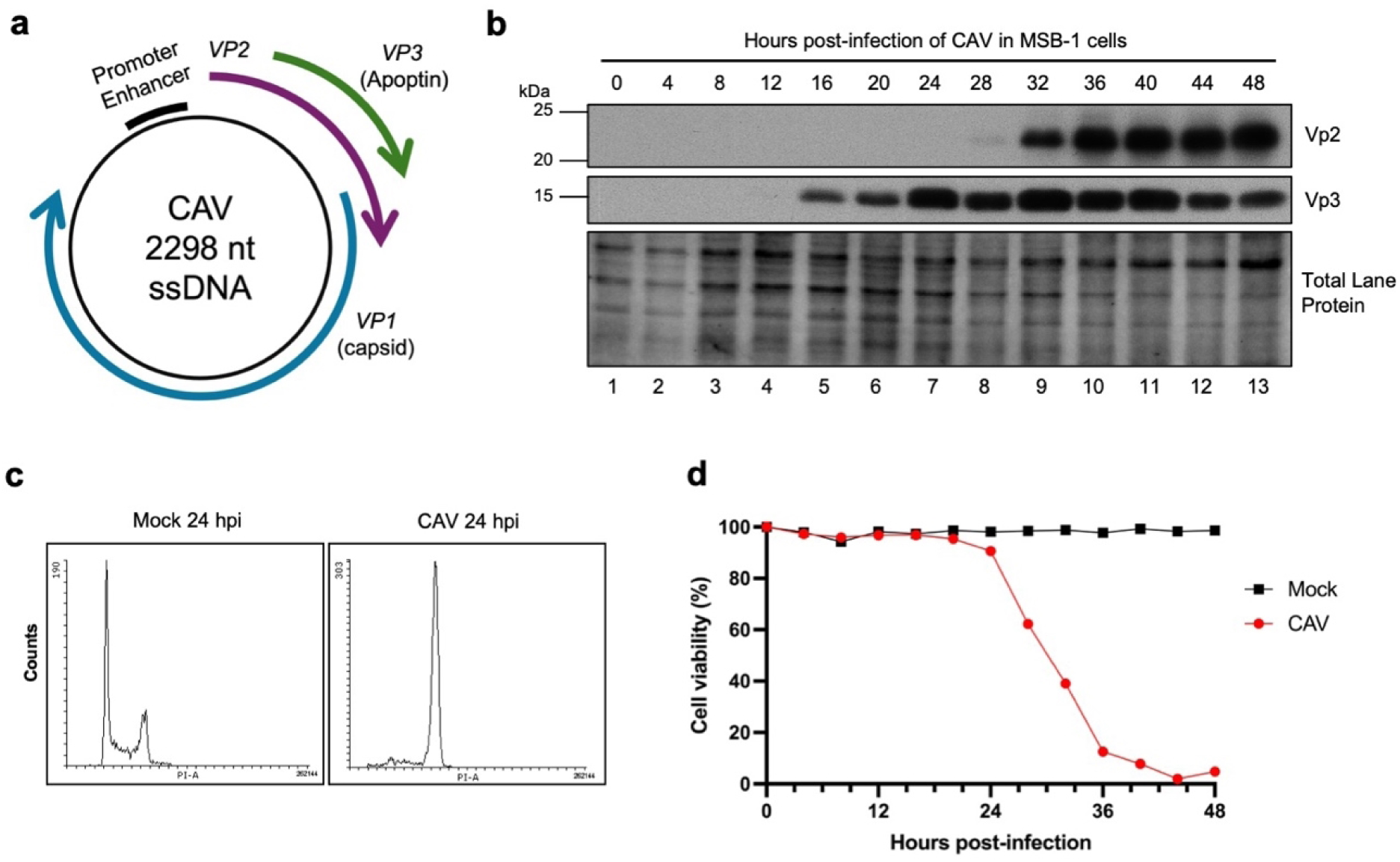
Viral protein expression in CAV-infected MSB-1 cells induces mitotic arrest and result in a cytopathic effect. (A) Schematic of the overlapping reading frames (ORFs) in the CAV genome coding for the three viral proteins of the virus. (B) Immunoblots showing the expression of Vp2 and Vp3 at 4-hour timepoints during CAV infection for 48 hours. Total lane protein is used as a loading control. (C) Flow cytometry analysis of mock– vs. CAV-infected cells stained with propidium iodide for DNA content at 24 hours post-infection (hpi). (D) Cell viability curves of mock-(black square) vs. CAV– (red circle) infected cells over the course of 48 hours. Cell viability was measured via trypan blue staining of infected cells every 4 hours. This figure is representative of three independent experiments.

## CAV infection results in mitotic arrest and a subsequent cytopathic effect

To determine how CAV affects cell cycle dynamics during a normal infection, we used an established infection model of CAV (17). Chicken MDCC MSB-1 cells were infected with CAV and monitored for viral gene expression, cell cycle state and induction of cell death (Fig 1). Fig 1B shows that the CAV Vp3 protein is expressed starting approximately 16 hours post-infection (hpi). Flow cytometry at 24 hpi shows that infection results in a near complete mitotic consistent with inhibition of the APC/C (Fig 1C). Starting at approximately 24 hpi, infected cells undergo a rapid cytopathic effect with all cells dead by 48 hpi (Fig 1D).

## Vp3 is important for the replication and cytopathogenicity of CAV

To further understand the role of viral APC/C binding proteins during replication, we derived a mutant CAV that was defective for expression of *VP3* (CAV Vp3-). As shown in the schematic in Fig 1A, the *VP3* open reading frame (orf) is completely contained in an alternative reading frame within the *VP2* replicase gene. Therefore, we derived CAV Vp3-such that the *VP3* initiation codon is mutated, while maintaining the *VP2* orf (Fig 2A). Fig 2B shows that viral replication of the CAV Vp3-mutant occurs at approximately one third of the wild-type CAV (CAV WT). Co-transfection of exogenous *VP3* with the CAV Vp3-replicative form (RF) partially rescued the replication deficiency of the mutant virus (Fig 2C). Loss of *VP3* expression was confirmed by immunoblot (Fig 2D). Interestingly, viral gene expression appeared to be delayed in the CAV Vp3-mutant with expression of the Vp2 replicase protein only becoming apparent 72 hpi compared to 48 hpi in the CAV WT. Moreover, the pronounced induction of mitotic arrest observed with the WT virus was absent in the CAV Vp3-, suggesting that the Vp3 protein is required for the cell cycle arrest (Fig 2E). Similarly, the cytopathic effect of the virus was also lost in the CAV Vp3-mutant (Fig 2F). Since the Vp3 protein binds and inactivates the APC/C complex (7), we then examined whether inhibition of the APC/C could rescue the defect in replication of CAV Vp3-. In Fig 2G and 2H, we show that addition of the APC/C inhibitor tosyl-L-Arginine Methyl Ester (TAME) can greatly rescue the replication defect caused by loss of Vp3. These data suggest that by interacting and inhibiting the APC/C, the Vp3 protein induces a cell state that is more amenable to viral replication.

**Fig 2.**
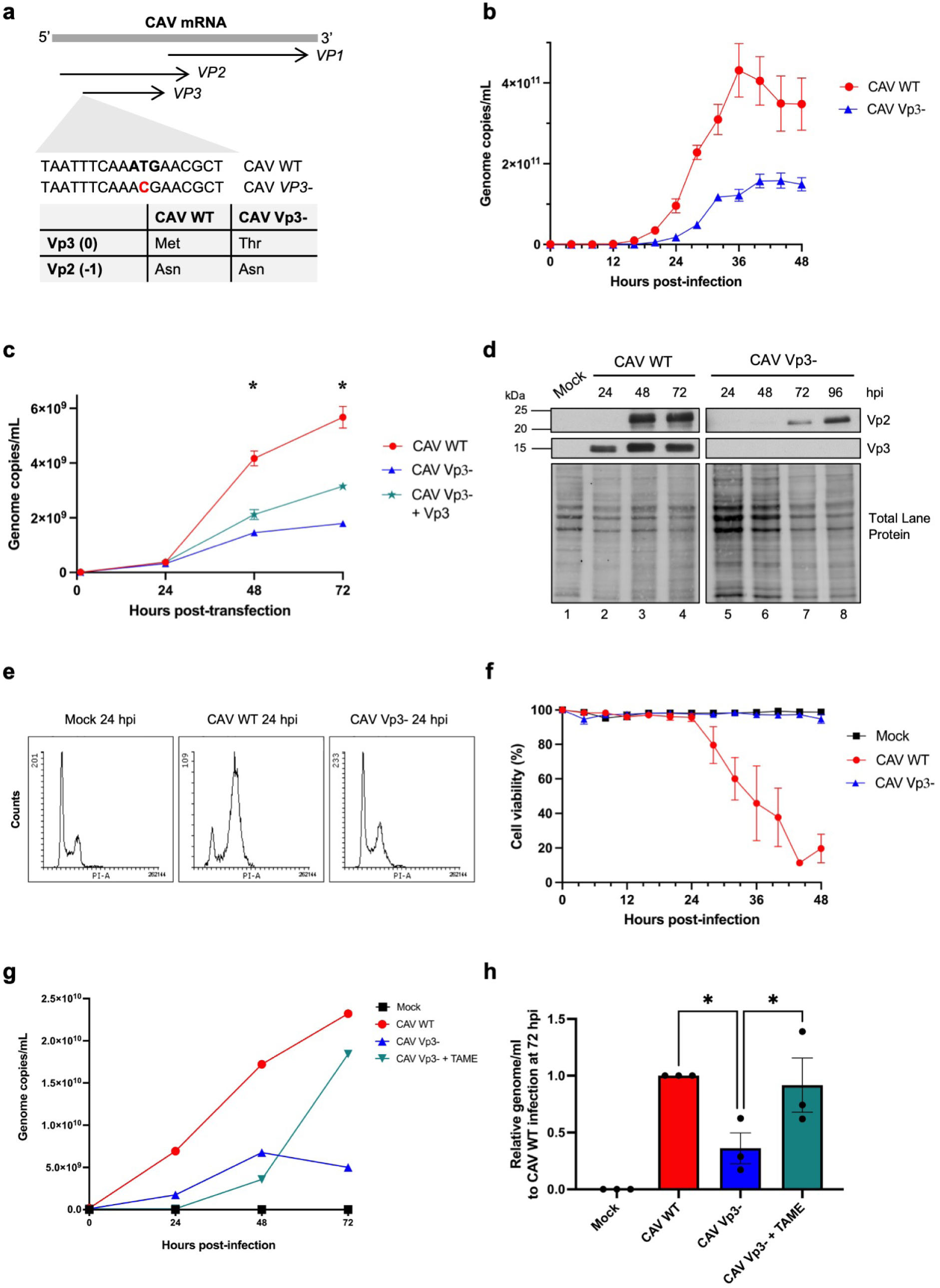
Vp3 is important for the replicative efficiency and cytopathogenicity of CAV. (A) Schematic of the mutagenesis strategy to generate the CAV Vp3-mutant construct. A T>C point mutation was made to abolish the start codon in the *VP3* reading frame. This also results in a silent mutation in the alternate reading frame for *VP2* and thus has no effect on the expression of *VP2*. (B) Viral growth curves for CAV WT (red circle) and CAV Vp3-(blue triangle) infected MSB-1 cells. Infected cells were taken every 4 hours for 48 hours and viral genomes were measured by qPCR at each timepoint. *N* = 3. (C) Quantification of viral genomes in MSB-1 cells transfected with CAV WT, CAV Vp3– or CAV Vp3-DNA with the p3XFLAG-VP3 expression vector. Transfected cells were taken every 24 hours for 72 hours and viral genomes were measured by qPCR at each timepoint. *N* = 3; *, *P* < 0.05 for all samples. (D) Immunoblots showing the temporal differences in Vp2 expression in the CAV WT vs. CAV Vp3-infections of MSB-1 cells. Hpi, hours post-infection. Total lane protein is used as the loading control. (E) Flow cytometry analysis of mock, CAV WT, and CAV Vp3-infected cells stained with propidium iodide for DNA content at 24 hours post-infection (hpi). (F) Cell viability curves of mock-(black square), CAV WT-(red circle) and CAV Vp3-(blue triangle) infected cells over the course of 48 hours. Cell viability was measured via trypan blue staining of infected cells every 4 hours. *N* = 3. (G) Viral growth curves for mock (black square), CAV WT (red circle) and CAV Vp3-(blue triangle) infected MSB-1 cells, as well as CAV Vp3-infected cells treated with TAME. Infected cells were taken every 24 hours for 48 hours and viral genomes were measured by qPCR at each timepoint. *N* = 3. (H) Quantification of viral genomes at the 72-hour timepoint of the infections in (G), relative to the CAV WT infection. *N* = 3; *, *P* < 0.05.

## Vp3 induces a metabolic environment that is favourable for nucleotide biosynthesis

We hypothesized that the Vp3 protein may enhance replication by inducing a more favourable metabolic state for viral replication. To test this hypothesis, metabolic profiling was performed on MSB-1 cells that were mock, CAV or CAV Vp3-infected. Metabolites of infected cells were extracted and quantified using liquid chromatography coupled to mass spectroscopy (LC-MS). Over 150 different metabolites were detected and quantitated (Tables S1A and S1B). Fig S1 shows a principal component analysis (PCA) plot of the metabolic changes observed in each of the replicate mock, CAV and CAV Vp3-samples. The PCA plot shows that there are major differences in metabolic profiles following infection with CAV. Mock infection and infection with the CAV Vp3-mutant clustered closer together, whereas the samples from WT-infected cells clustered tightly within a discrete outlying area of the plot. In Fig 3A, pathway analysis was performed by using the MetaboAnalyst software to identify the most affected metabolic pathways during CAV infection. Intriguingly, pyrimidine and purine metabolism were two of the most significantly impacted pathways following infection (Fig 3A and Table S2). The heat map in Fig 3B shows that a wide range of intermediate metabolites in pyrimidine and purine metabolism were elevated in CAV-infected cells but not in the CAV Vp3-infection. These data demonstrate a shift in host cell nucleic acid metabolism during CAV infection to facilitate viral genome replication.

**Fig 3.**
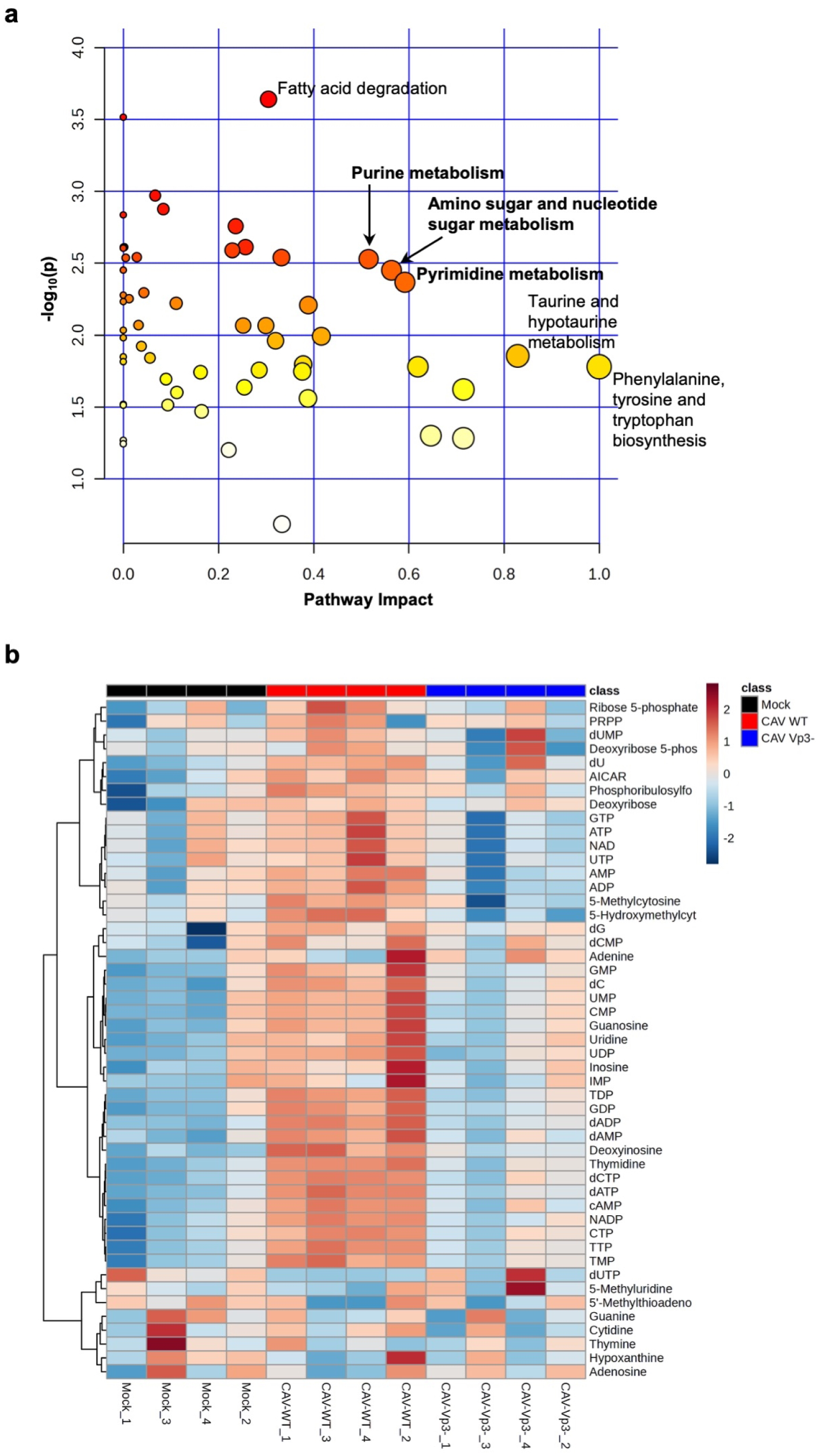
Vp3 induces a metabolic state that is more conducive to nucleotide metabolism. (A) Pathways analysis from comparing metabolites of MSB-1 cells infected with CAV WT relative to that of CAV Vp3-at 24 hours. The metabolites of the infected cellular extracts were analyzed by LC-MS. Raw metabolite data were analyzed and presented using the MetaboAnalyst web-based software. Pathway importance is a combination of pathway impact score (the higher score, the higher the biological impact of the pathway – *x axis*) and significance (the higher the – log_10_(*P*) value, the more statistically significant those changes are for the pathway – *y* axis). The colour of each circle is based on the enrichment *P* value (white-red gradient), while the size of each plot represents the impact value. *N=4.* (B) Heatmap visualizing the changes in all nucleotides and derivatives detected by LC-MS in the CAV WT and CAV Vp3-infections from (A), as well as mock infection. Values are colour-coded on a blue-red gradient and are centred around 0, at which no relative change in levels is detected.

## APC/C inhibition by Vp3 elevates the expression of APC/C substrates involved in nucleotide biogenesis

The APC/C regulates mitotic progression by targeting substrates for degradation in the form of APC/C^CDC20^. Inhibition of the APC/C by Vp3, as demonstrated in the aforementioned experiments, explains the resulting mitotic arrest and cytopathogenicity from CAV infection. However, in addition to its mitotic functions, the APC/C complexed with CDH1 controls S-phase entry by targeting a variety of substrates including nucleotide biosynthetic enzymes. At the G1-S boundary, APC/C^CDH1^ is inactivated allowing the stabilization of enzymes including ribonucleotide reductase regulatory subunit M2 (RRM2)(24), thymidine kinase 1 (TK1)(25), and thymidylate kinase (TMPK)(26). We therefore hypothesized that viral targeting of the APC/C may be a mechanism to stabilize enzymes required for nucleotide biosynthesis. To test this possibility, we examined the expression of RRM2 in MSB-1 cells infected with CAV. Fig 4A shows that at 24 hpi expression, the protein levels of RRM2 are elevated compared to the mock infection or the CAV Vp3-infection. A time course of infection showed that RRM2 levels become elevated as Vp3 expression becomes detectable at approximately 12 hpi (Fig 4B). Fig 4C shows that the concentrations of deoxythymidine monophosphate (dTMP), deoxythymidine diphosphate (dTDP) and deoxythymidine triphosphate (dTTP) were all significantly elevated following CAV infection but were not affected in the CAV Vp3^-^ infections. A similar trend was also observed for the other deoxynucleotide triphosphates (dNTPs). Despite the elevation in dNTPs, no significant changes were observed in the levels of the nitrogenous bases suggesting that these effects are due to enhanced salvage pathways rather than *de novo* biosynthesis (Fig S2). Since we previously observed that the Vp3 protein can inhibit the APC/C when expressed alone (7), we determined if it can also stimulate nucleotide biosynthesis outside the context of a CAV infection. Fig 4D shows that when Vp3 is expressed in H1299 cells, there is an increase in the steady state levels of RRM2 and TK1. Importantly, the levels of the mRNAs encoding these enzymes were not affected in infected cells, suggesting that the elevated expression was due to increased protein stability (Fig S3). Finally, we show that expression of Vp3 alone was able to significantly elevate production of dTMP, dTDP and dTTP (Fig 4E), indicating that the protein is both necessary and sufficient to elevate nucleotide levels. Taken together, these data show that viral targeting of the APC/C is a mechanism to elevate cellular biosynthesis of nucleotides and maximize viral replication.

**Fig 4.**
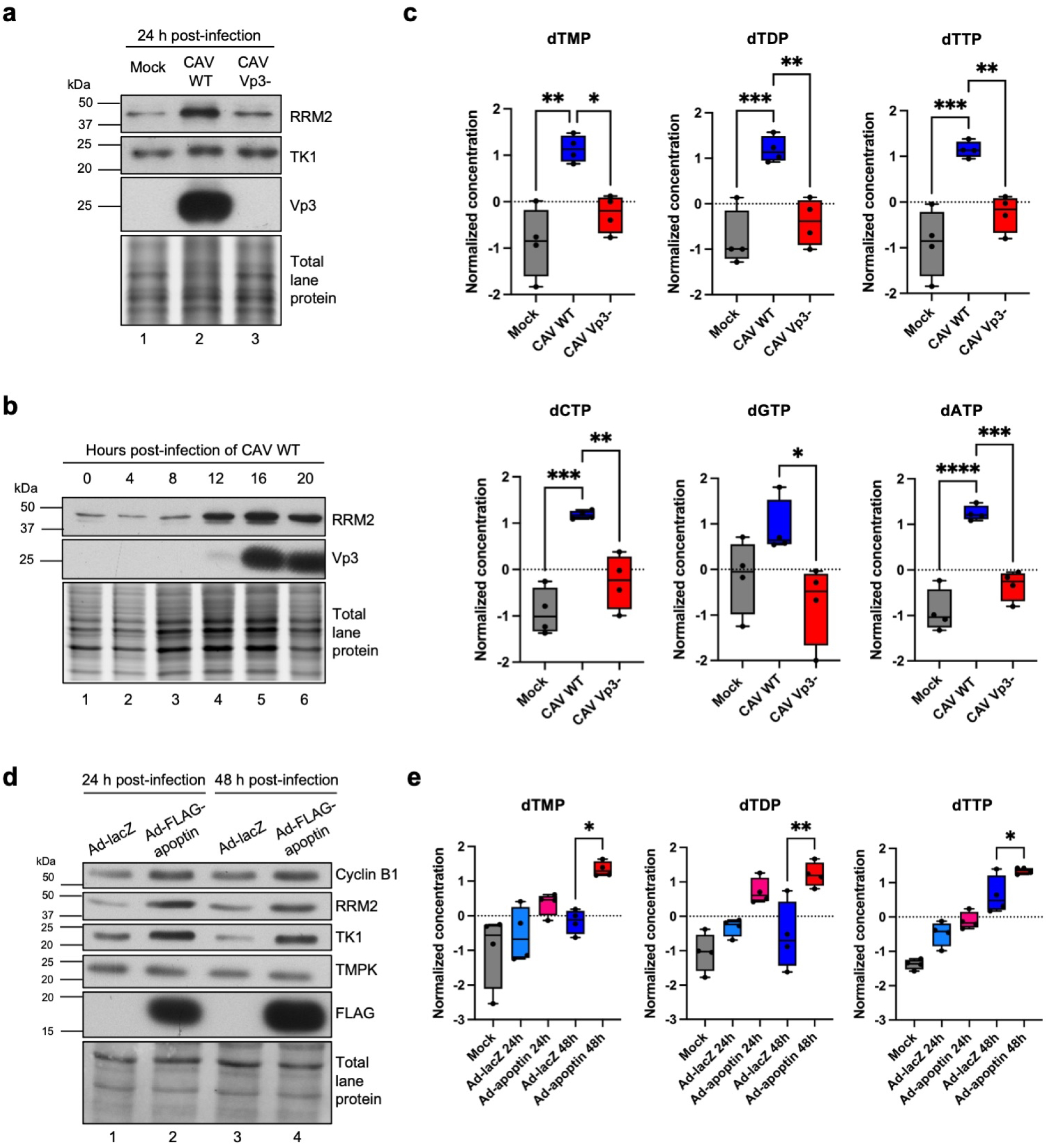
Vp3 is necessary and sufficient to increase nucleotide synthesis by elevating levels of APC/C substrates that are involved in nucleotide biosynthesis. (A) Immunoblots comparing the expression of RRM2 and TK1, both substrates of the APC/C, in lysates of MSB-1 cells at 24 hours post-infection with mock, CAV WT or CAV Vp3-. Total lane protein is used as the loading control. (B) Immunoblots tracking the temporal changes in RRM2 expression in MSB-1 cells during the initial hours of CAV WT infection. Timepoints were taken every 4 hours for the first 20 hours of CAV infection. Total lane protein is used as the loading control. (C) Normalized levels of dTTP, its precursors and the other dNTPs compared across the different infection conditions from (A). *N* = 4. (D) Immunoblots showing the expression of APC/C substrates involved in nucleotide biosynthesis in H1299 cells overexpressing the Vp3/apoptin protein via the FLAG-tagged apoptin adenovirus (Ad-apoptin). The Ad-lacZ virus was used in the control infections. Infected cells were harvested at the 24– and 48-hour timepoints post-infection. Cyclin B1 is used as a mitosis marker. Total lane protein is used as the loading control. (E) Normalized levels of dTTP and its precursors compared across the different infection conditions from (D). *N* = 4. *, *P* < 0.05; **, *P* < 0.01; ***, *P* < 0.001, ****, *P* < 0.0001.

## Discussion

Viruses have relatively small genomes, necessitating the evolution of highly efficient mechanisms to exert changes in cellular physiology during replication. In the current study, we used CAV to study the impact of viral APC/C inhibition on viral replication. Our findings demonstrate the importance of viral inhibition of the APC/C in upregulating nucleotide biosynthesis to facilitate viral replication. This mechanistic finding can be extended to a diverse range of viruses to explain why they have convergently evolved to inhibit the APC/C. CAV is one of the smallest viruses that causes pathology in animals with a genome of only 2.3 kilobases (15). Despite encoding only three polypeptides, the virus has maintained the inhibitory interaction with the APC/C, underscoring the importance of targeting this complex. The CAV Vp3 protein induces the disassembly of the APC/C by binding the large scaffold subunit of the APC/C called APC1 (7). Similar changes are observed during infections with human cytomegalovirus (HCMV), a betaherpesvirus (4, 27–29), and HIV, whose VPR protein also binds APC1 and stimulates its degradation (6). Besides binding to the APC1 scaffold protein, other viral APC/C inhibitory mechanisms have been reported. The adenovirus E4orf4 protein was shown to recruit the PP2A phosphatase to the APC/C resulting in the dephosphorylation and inactivation of the complex (3) and the human papillomavirus (HPV) E2 protein was shown to bind and sequester both the CDH1 and CDC20 co-activators of the APC/C (2). Perhaps the most dramatic example of viral APC/C inhibition is observed in some types of poxviruses. Orf virus (ORFV) is a poxvirus that encodes a viral homolog of the catalytic APC11 subunit of the APC/C termed the poxvirus APC/cyclosome regulator (PACR)(5). The PACR protein is catalytically inert and displaces the endogenous APC11 resulting in an inactive APC/C (30). In all these studies, viruses appear to inactivate both the APC/C^CDH1^ and the APC/C^CDC20^ complexes. These two complexes have very different effects on cell cycle progression. APC/C^CDH1^ has an inhibitory effect on the cell cycle progression by elevating levels of CDK inhibitors such as p21, p57 and p27 (31, 32). APC/C^CDH1^ inhibition and transition to S phase in CAV-infected cells would be needed for the elevation of nucleotide pools to facilitate viral genome replication. In contrast, APC/C^CDC20^ is absolutely required for mitotic progression and as observed in CAV-infected cells, inhibition of both complexes would be predicted to result in mitotic arrest.

Despite the numerous studies showing viral inhibition with the APC/C, a molecular understanding of the role of these interactions during infection has never been shown. Although APC/C inhibition may enhance viral replication through more than one mechanism, the metabolomics data presented in this study suggest that upregulation of nucleotide biosynthesis is a key benefit of virally-induced APC/C inhibition. Other evidence also points to the importance of APC/C inhibition to elevate nucleotide pools. For example, poxviruses either encode a copy of PACR, or a viral thymidine kinase but never both, suggesting that these two proteins have redundant functions in stimulating nucleotide production (5, 33). Similarly, although most herpesviruses encode a copy of thymidine kinase, HCMV, which is known to inhibit the APC/C with its pUL97 and pUL21 proteins during infection, does not (4, 27–29). The work presented here defines a previously unappreciated importance for APC/C inhibition that is broadly applicable to a wide range of viruses.

## Methods and Methods

### Cell lines and maintenance of cell culture

Marek’s Disease chicken cell (MDCC) MSB-1 cells (CLS Cell Lines Service), a chicken lymphoblastoid cell line immortalized by Marek’s Disease Virus (MDV), were cultured in RPMI-1640 (Wisent Bioproducts) supplemented with 10% fetal bovine serum (Sigma, Canadian origin) and 50 μg/mL gentamycin (Wisent Bioproducts). Cells were kept at 41°C under 5% CO_2_ atmosphere. MSB-1 cells were split every 2 days and maintained at a density of 0.3–1.5 × 10^6^ cells/mL. H1299 cells (ATCC) were cultured using DMEM (Wisent Bioproducts) with 10% FBS and were split every 2-3 days. H1299 cells were maintained at 37°C under 5% CO_2_ atmosphere.

### Plasmids

The full-length Cux-1 strain CAV genome (Genbank accession number NC_001427, from Dr. Mathieu Noteborn) was ligated into the pIC20H expression plasmid and was amplified in DH5α *Escherichia coli.* The CAV VP3-mutant construct was made by introducing a single point mutation in the ATG start codon of the *VP3* ORF via site-directed mutagenesis (see **Fig. 2a**): here, the T>C nucleotide change results in a missense mutation (Met to Thr) that eliminates the start codon of the *VP3* ORF, while the coding sequence for *VP2* remains intact as it is a silent mutation (Asn to Asn) in the overlapping *VP2* ORF. VP3 (apoptin) cDNA was cloned into the p3XFLAG-*myc*-CMV-26 expression vector (Sigma).

### Replicative form (RF) viral DNA preparation, transfection, and virus production

To obtain the replicative form (RF) of CAV for virus production, the purified pIC20H CAV WT or VP3-plasmid was linearized by EcoRI and BglI. The linearized CAV genome was purified by agarose gel and re-circularized with T4 DNA ligase (NEB) at a low concentration (1 ng/mL) to maximize intramolecular ligation. The circular, RF CAV was then recovered using a silica spin column, following the PCR clean-up instructions from the GenepHlow Gel/PCR Kit (Geneaid). Circular RF CAV was confirmed by agarose gel before transfection.

For virus production, 400–900 ng of circular RF CAV WT or CAV VP3-were electroporated into 2 × 10^6^ MSB-1 cells (seeded at a density of 3 × 10^5^ cells/mL the day before) using the AMAXA Nucleofector II device (custom T-001 setting) and Reagent Kit T (Lonza). Following electroporation, cells were resuspended in growth media at 3 × 10^5^ cells/mL to recover for 1 hour at 41°C and washed with phosphate buffered saline (PBS) to remove residual RF CAV. Cells were grown for 3 days post-transfection, and 140 μL of the cell suspension was taken every 24 hours (or otherwise specified) for viral genome quantification. After 72 hours, cells were lysed by three freeze-thaw cycles and the liberated virus in the media was filtered through a 0.45-μm membrane (Corning) to collect the cell-free viral supernatant. A final 140-μL sample was taken for viral genome quantification. Viral supernatants were stored at –80°C until use for infection.

### Viral genome quantification

Viral DNA was extracted from the 140-μL samples of virus-containing MSB-1 cells and filtered cell-free viral supernatants using the QIAamp Viral RNA Mini Kit (QIAGEN) according to the manufacturer’s instructions. Extracted DNA was digested with DpnI remove traces of bacterial RF CAV from the initial electroporation step. Viral genomes were quantified by qPCR based on a standard curve generated by known dilutions of pIC20H-CAV DNA, using the SsoAdvanced Universal SYBR Green Supermix (Bio-Rad) and the CFX Connect Real-Time PCR System (Bio-Rad). Primers for qPCR targeted the *VP1* gene (common to both the CAV WT and CAV Vp3-) in such a way that the resulting amplicon spanned a DpnI restriction site. The primer sequences used are as follows: forward – ATGACCCTGCAAGACATGGG; reverse – CTTTTTGCCACCGGTTCTGG. Following viral genome quantification, the concentrations of both CAV WT and Vp3-supernatants were equalized by the addition of fresh media.

### CAV infection of MSB-1 cells

MDCC MSB-1 cells were seeded at 3 × 10^5^ cells/mL one day prior to infection. Five-mL aliquots of virus-laden media of at least 1 × 10^9^ viral genomes/mL were thawed and used to resuspend 10 million MSB-1 cells for each infection. Cells were incubated with WT or Vp3-virus (or media for mock infection) for 1 hour at 41°C. The viral suspension was then removed, and the infected cells were washed with PBS. Finally, the infected cells were resuspended at 3 × 10^5^ cells/mL with regular growth media. Cell suspension aliquots of 140 μL of each infection were taken at specific timepoints post-infection where needed and viral genomes were quantified as described above.

### Cell counts and viability

At specific timepoints post-infection, cells of each infection were counted using a hemocytometer and percent cell viability was determined by Trypan Blue staining (1:10, dye: cells) under light microscopy.

### Cell cycle analysis (flow cytometry)

Cells of each infection were collected and fixed with ice-cold ethanol. Cells were stained with propidium iodide (PI, from Sigma) solution (100 ug/mL PI, 0.2 mg/mL RNase A, 0.6% Nonident P-40 in PBS). Cellular DNA was analyzed using the LSRFortessa (BD Biosciences). Data were analyzed with Flowing Software 2.5.1.

### Western blotting

Infected MSB-1 cells taken at different timepoints post-infection were lysed in 1× Laemmli buffer (diluted in ddH_2_O from a 4× stock: 200 mM Tris-HCl pH 6.8, 4% SDS, 40% glycerol, 4% β-mercaptoethanol, 0.012% bromophenol blue, and 50 mM DTT). Cell pellets were sonicated, and DNA was further sheared using a 23-gauge needle (BD PrecisionGlide) and a 1-cc syringe (BD). Twenty micrograms of total protein were separated by SDS-PAGE and probed with antibodies against Vp2 (custom made), Vp3 (Dr. Mahvash Tavassoli Lab), RRM2 (Invitrogen), TK1 (Santa Cruz), Cyclin B1 (Santa Cruz), FLAG (Sigma), and TMPK (Dr. Zee-Fan Chang Lab). Total lane protein was used as the loading control: polyacrylamide gels with 0.5% (by volume) of 2,2,2-trichloroethanol (TCE, from Sigma) were used to resolve total protein, after which the gel was activated using the ChemiDoc XRS+ stain-free enabled UV transilluminator (Bio-Rad) with Image Lab 6.0.1 (Bio-Rad) to visualize total lane protein. Total protein transferred onto PVDF membranes was also visualized with the ChemiDoc XRS+ UV transilluminator.

### Vp3-rescue experiments

For the Vp3 rescue experiment, 1 μg of the p3XFLAG-*myc*-CMV-26 plasmid expressing exogenous *VP3* was co-transfected with the circular RF CAV *VP3*-genome as described above. For the TAME rescue experiment, CAV Vp3-virus was used to infected MSB-1 cells as described above. Immediately following infection, pro-N-4-tosyl-L-arginine methyl ester (proTAME) (Boston Biochem) in DMSO or DMSO only (vehicle) was added to culture media to a final concentration of 10 μM. For both rescue experiments, aliquots of cell suspensions were taken for quantification as described above.

### Cell extraction for metabolite analysis

All LC-MS grade solvents and salts were purchased from Fisher (Ottawa, Ontario Canada: water, acetonitrile (ACN), Methanol (MeOH), formic acid, ammonium acetate and ammonium formate. The authentic metabolite standards and N-ethylmaleimide were purchase from Sigma-Aldrich Co. (Oakville, Ontario, Canada).

Cells were washed with ice-cold 150 mM ammonium formate solution pH of 7.4 containing 1 mg/mL N-ethylmaleimide to protected free thiols (34). Cells were then scraped and extracted with 600 μL of 31.6% MeOH/36.3% ACN in H_2_O (v/v also containing 1 mg/mL N-ethylmaleimide). Cells were lysed and homogenized by bead-beating for 30 seconds at 50 Hz using 4 ceramic beads (2 mm) per sample (TissueLyser II – Qiagen). Cellular extracts were partitioned into aqueous and organic layers following dimethyl chloride treatment and centrifugation. The aqueous supernatants were dried by vacuum centrifugation with sample temperature maintained at −4°C (Labconco, Kansas City MO, USA). Dried extracts were subsequently re-suspended in 50 μL of chilled H_2_O and clarified by centrifugation at 1°C. Sample injection volumes for analyses were 5 μL per injection.

### LC-MS

For targeted metabolite analysis, samples were injected onto an Agilent 6470 Triple Quadrupole (QQQ)–LC–MS/MS (Agilent Technologies). Chromatographic separation of metabolites was achieved by using a 1290 Infinity ultra-performance quaternary pump liquid chromatography system (Agilent Technologies). The mass spectrometer was equipped with a Jet Stream^TM^ electrospray ionization source, and samples were analyzed in negative mode. The source-gas temperature and flow were set at 150 °C and 13 L min^−1^, respectively, the nebulizer pressure was set at 45 psi, and capillary voltage was set at 2,000 V. Multiple reaction monitoring parameters (qualifier/quantifier ions and retention times) were either obtained optimized using authentic metabolite standards.

Chromatographic separation of the isomers and other metabolites was achieved by using a Zorbax Extend C18 column 1.8 μm, 2.1 × 150mm^2^ with guard column 1.8 μm, 2.1 × 5mm^2^ (Agilent Technologies). The chromatographic gradient started at 100% mobile phase A (97% water, 3% methanol, 10 mM tributylamine, 15 mM acetic acid, 5 µM medronic acid) for 2.5 min, followed with a 5-min gradient to 20% mobile phase C (methanol, 10 mM tributylamine, 15 mM acetic acid, 5 µM medronic acid), a 5.5-min gradient to 45% C and a 7-min gradient to 99% C at a flow rate of 0.25 mL min^−1^. This was followed by a 4-min hold time at 100% mobile phase C. The column was restored by washing with 99% mobile phase D (90% ACN) for 3 min at 0.25 mL min^−1^, followed by increase of the flow rate to 0.8 mL min^−1^ over 0.5 min and a 3.85-min hold, after which the flow rate was decreased to 0.6 mL min^−1^ over 0.15 min. The column was then re-equilibrated at 100% A over 0.75 min, during which the flow rate was decreased to 0.4 mL min^−1^, and held for 7.65 min. One minute before the next injection, the flow was brought back to forward flow at 0.25 mL min^−1^. For all LC–MS analyses, 5 μL of sample was injected. The column temperature was maintained at 35°C. Relative concentrations were determined from external calibration curves. Data were analyzed by using MassHunter Quantitative Data Analysis B.10.00 (Agilent Technologies). Data presented are peak area normalized by cell counts at the time of the cell extractions.

### Analysis of metabolites

Data were analyzed and presented using the web-based platform MetaboAnalyst (35). The one-factor Statistical Analysis module was used to generate the PCA plot of all infections. Data were normalized with log-transformation and auto-scaling before analysis. To compare the individual metabolites from the different infections, a one-way ANOVA with post-*hoc* Tukey’s HSD multiple comparisons tests were performed with a *P* value cutoff of 0.05. Next, the Pathways Analysis module was used to determine the significantly impacted molecular pathways in the CAV WT infections relative to the CAV Vp3-infections. Data were normalized with log-transformation and auto-scaling before analysis. The following pathway analysis parameters were selected: Scatter plot (Visualization method), Global Test (Enrichment method), Relative-betweeness Centrality (Topology measure). The *G. gallus* (KEGG) pathway library provided by the platform was selected as the Reference metabolome. To generate the heatmap comparing nucleotide changes, the one-factor Statistical Analysis module was used to analyze only nucleotides and their derivatives across all infection conditions. Data were normalized with log-transformation and auto-scaling before analysis. The following parameters were used to display the normalized data: Autoscale features (Standardization), Euclidean (Distance measure), Ward (Clustering method).

### Overexpression of apoptin in H1299 cells

To express apoptin in mammalian cells, the FLAG-tagged apoptin adenovirus (Ad-apoptin) was used as described previously (7). The Ad-lacZ virus was used as the control. Both Ad-apoptin and Ad-lacZ viruses were generated and titered as described previously (7). H1299 cells were seeded at around 60% confluency and were infected at an MOI of 35. Cells were harvested at the relevant timepoints and were processed for Western blot and metabolite analyses as described above.

### Gene expression analysis by RT-qPCR

Total RNA was extracted from MSB-1 cells using the RNesay Mini Kit (Qiagen). Nine hundred nanograms of total RNA were reverse transcribed into cDNA using the QuantiTect Reverse Transcription Kit (Qiagen). qPCR was performed using SsoAdvanced SYBR Green Supermix (Bio-Rad) with the CFX Real-Time system (Bio-Rad). All the reactions were performed in duplicate (technical). *ACTB* and *GAPDH* were used as reference genes for relative gene expression analysis via the 2^−ΔΔCT^ method. All primers were validated before assaying relative gene expression. qPCR primer sequences are listed as follows:

*RRM2*: GCATTTGCAGCAGTGGAAGG & TCACAGTGCAAACCCTCATC

*TK1*: GAGGGGCAGTTTTTCCCAG & GAGGATGCTCCCAAAAGCC

*DTYMK* (TMPK): GCTGGGAGCACGTAACATTG & ACTTGTGAAGGCCACTCCAG

*ANAPC1*: CACCCAGAGAACCTTTGCC & GCAGAGCCAAACACACAGAC

*ANAPC4*: GCTTCTGGGTGATGTAAGGC & CAAGAACCTGCCACCTCAG

*GAPDH*: CCCTGAGCTGAATGGGAAGC & TCAGCAGCAGCCTTCACTAC

*ACTB*: CGTTACTCCCACAGCCAGC & GGGCGACCCACGATAGATG

### Statistical analysis

Statistical analysis was performed with the GraphPad Prism (Version 10) software (GraphPad) or on the web-based MetaboAnalyst platform (35). For comparisons of more than two groups, an ordinary one-way ANOVA was performed followed by *post hoc* pairwise comparisons (Dunnett or Tukey multiple comparisons test). A *P* value <0.05 is considered statistically significant. For histograms and curves, data are expressed as mean ± SEM. For box & whisker plots, all data points are shown with the median, maximum and minimum. Biological replicates were used to generate the plots in the figures. A biological replicate for a given condition is defined by an independent sample measured in an experiment or a sample measured in an independently repeated experiment.

### Data and Code Availability Statements

All acquired raw metabolomics data were processed, normalized and analysed by the web-based MetaboAnalyst platform (35). For more information on how the data were analysed, see the “Analysis of metabolites” section.

## Acknowledgements

We thank the support of the Metabolomics Innovation Resource Core at the Goodman Cancer Institute for the assistance in experimental design and data collection. We also acknowledge the support from the McGill Flow Cytometry Innovation Platform. The Vp3 and TMPK antibodies were gifts from Dr. Mahvash Tavassoli and Dr. Zee-Fan Chang, respectively. O.J. Chen is a recipient of the Doctoral Training Scholarship from the *Fonds de recherche du Québec – Santé* (FRQS) and the Canada Graduate Scholarship – Doctoral (CGS-D) from the Canadian Institutes of Health Research (CIHR). C.Y. Feng is a recipient of the Canada Graduate Scholarship – Master’s (CGS-M). This work is funded by a project grant from the CIHR to J.G. Teodoro, and a Discovery Grant from the Natural Sciences and Engineering Research Council of Canada (NSERC) to J.G. Teodoro.

## Authors’ Contributions

O.J.C. and C.Y.F. contributed equally to this work, designing and carrying out most experiments. O.J.C. performed the metabolomics studies and the western blot experiments. C.Y.F. performed the viral replication and flow cytometry studies. R.J.L. assisted in collecting time points for the western blot and gene quantification studies. M.A.C. helped with the metabolomics studies. M.S.T.R. collected the raw metabolomics data. Y.W. quantified the gene expression via RT-qPCR. P.B. helped with the amplification and titration of the adenoviruses. Y.L. assisted with the viral replication experiments. D.S. established the viral replication studies. M.M.-K. generated the Vp3/apoptin expression vector. I.G. provided technical assistance with the viral replication and quantification protocols. N.S. provided supervision. D.A. provided guidance in the design of the metabolomics studies and finalized the raw metabolomics data. O.J.C., C.Y.F., D.S. and J.G.T. conceived the project. O.J.C. and C.Y.F. analysed the data. J.G.T. and O.J.C. wrote and revised the manuscript, respectively. All authors discussed the results and manuscript. J.G.T. supervised the project.

## Supporting Information

**Table S1A**: Raw data of metabolites of mock, CAV WT and CAV Vp3-infections in MSB-1 cells detected by LC-MS

**Table S1B**: Normalized metabolite data of mock, CAV WT and CAV Vp3-infections

**Table S2**: Significance and impact scores for the pathways analysis of metabolites from MSB-1 cells infected with CAV WT vs. CAV Vp3-

**Fig S1**: Principal component analysis (PCA) plot of the metabolic profiles of mock, CAV WT and CAV Vp3-infected MSB-1 cells at 24 hours post-infection

**Fig S2**: Normalized levels of nitrogenous bases compared across mock, CAV WT and CAV Vp3-infections

**Fig S3**: mRNA transcript levels of different enzymes involved in nucleotide synthesis during CAV infection

## Figures and Legends

**Fig S1.**
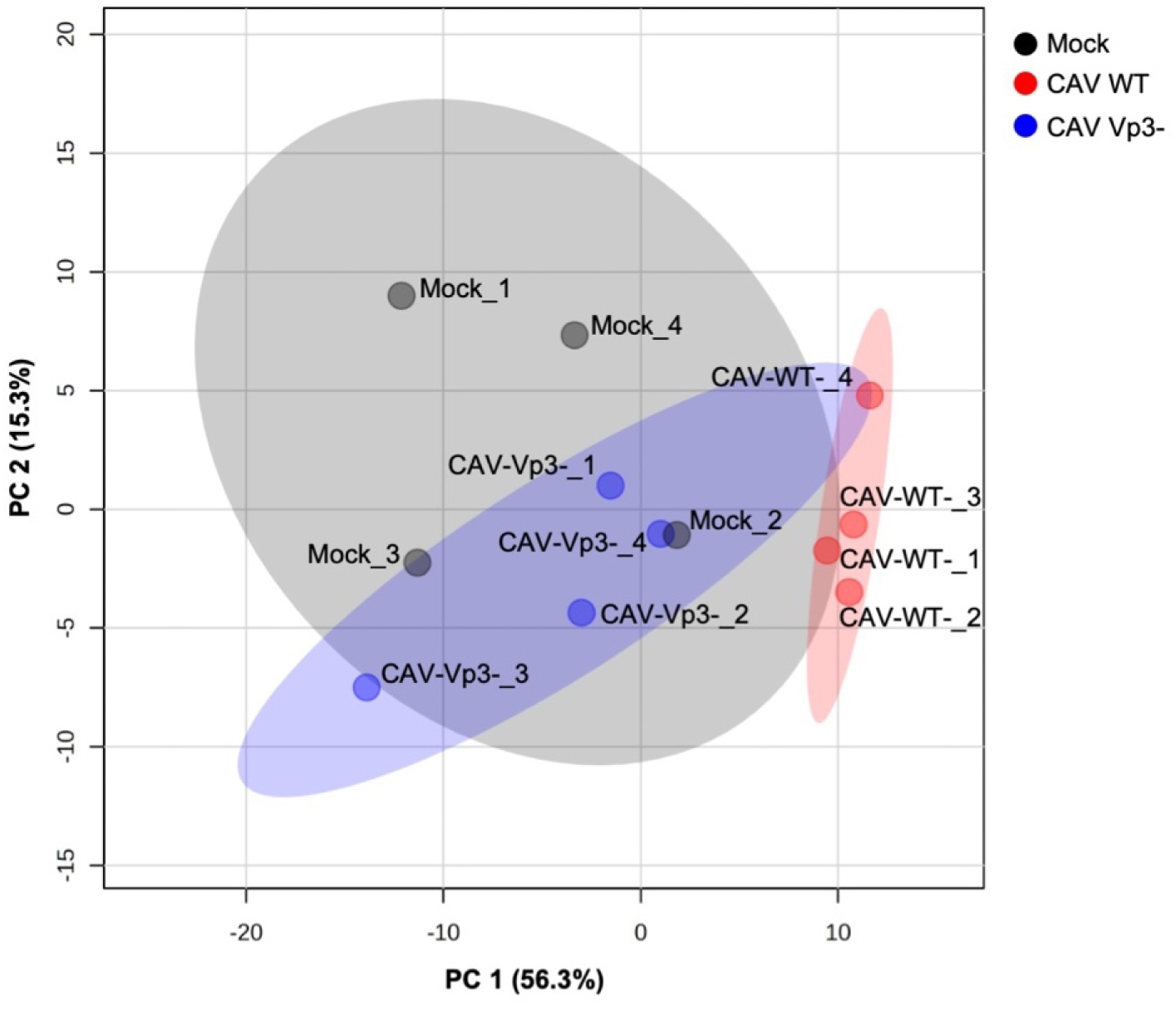
Principal component analysis (PCA) plot of the metabolic profiles of mock, CAV WT and CAV Vp3-infected MSB-1 cells at 24 hours post-infection.

**Fig S2.**
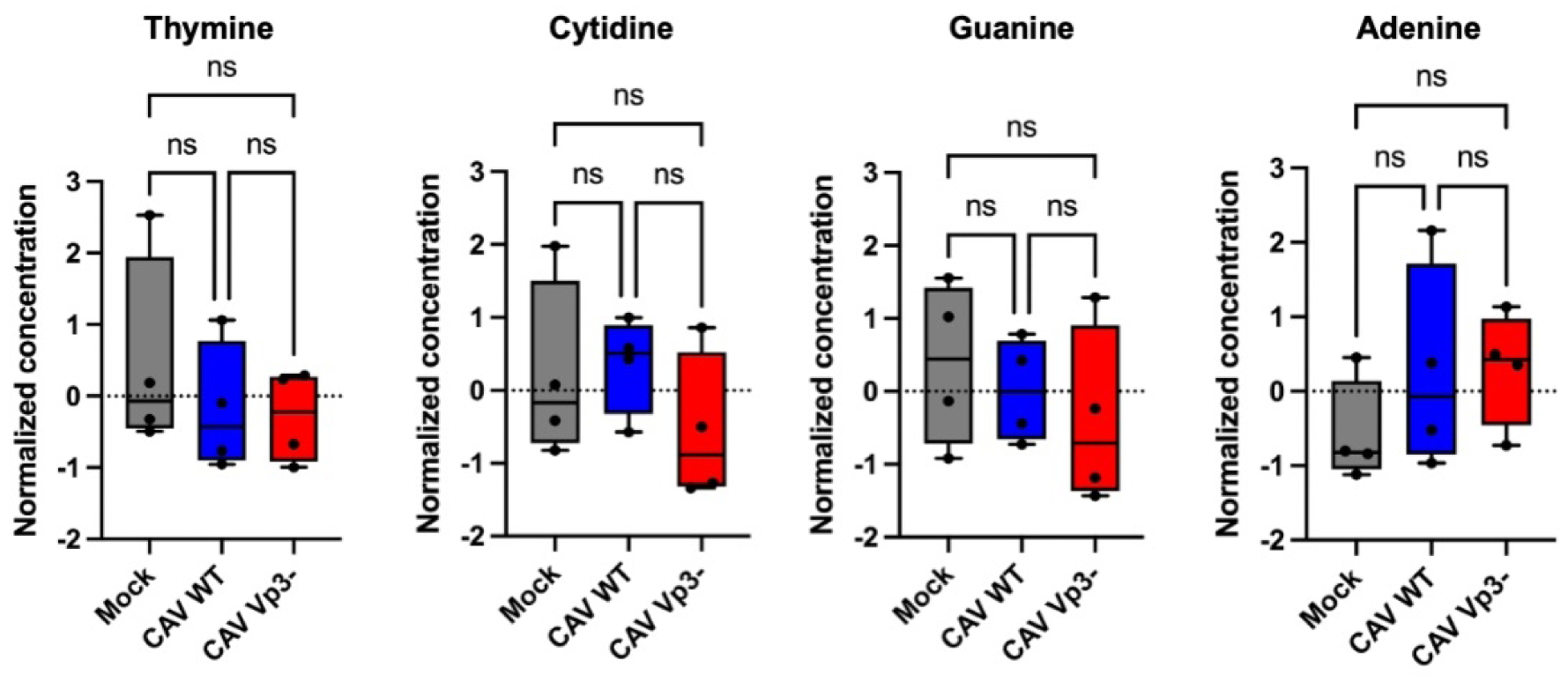
Normalized levels of nitrogenous bases compared across mock, CAV WT and CAV Vp3-infections. *N* = 4; ns, not significant.

**Fig S3.**
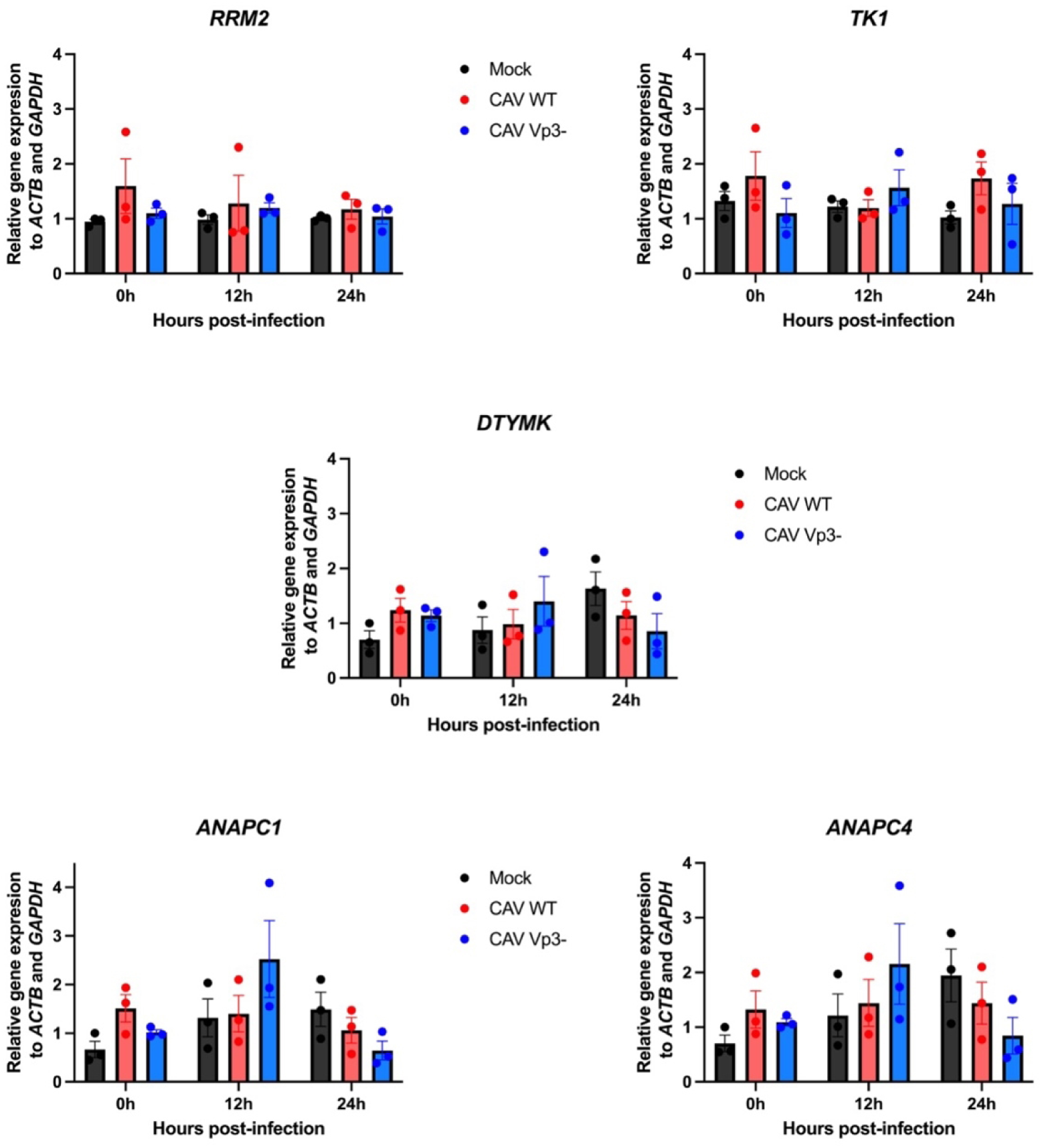
mRNA transcript levels of different enzymes involved in nucleotide synthesis during CAV infection. Relative transcript levels of *RRM2*, *TK1* and *DTYMK* (TMPK) in cells infected with mock, CAV WT or CAV Vp3-at 12-hour timepoints were measured by RT-qPCR. Gene expression was normalized to that of reference genes *ACTB* (actin) and *GAPDH*. *ANAPC1* (APC1) and *ANAPC4* (APC4) were used as controls for comparison. Each gene at every timepoint was measured from three independent experiments. *N* = 3 for all groups. Statistical significance for each comparison: n.s. (not significant; *P* > 0.05).

